# MetaScope: A High-Resolution Framework for Species-Level 16S Metataxonomic Classification

**DOI:** 10.1101/2025.08.29.673160

**Authors:** Sean Lu, Aubrey R Odom, Avi Shah, Yaoan Leng, Kiloni Quiles, W Evan Johnson

## Abstract

Accurate species-level classification of microbial communities remains a major challenge in microbiome analysis, particularly when using traditional 16S rRNA amplicon sequencing pipelines such as QIIME2 and DADA2. These methods often fail to resolve taxonomy beyond the genus level due to limitations in clustering algorithms and ambiguous marker genes. To address this, we present MetaScope, a modular, R-based software package that reimplements and extends the PathoScope 2.0 framework for high-resolution microbial classification. MetaScope introduces two key innovations: (1) the integration of user-defined or empirical prior weights into the Bayesian read reassignment algorithm to improve abundance estimation, and (2) MetaBlast, a secondary BLAST-based validation module for refining species-level assignments. Benchmarking against mock and clinical datasets demonstrates that MetaScope significantly outperforms QIIME2 and DADA2, achieving up to 93.6% species-level classification accuracy. This translates to offering enhanced resolution for downstream analyses such as ecological diversity metrics. These results highlight MetaScope as a powerful tool for advancing microbial community profiling in both research and clinical settings.

## Introduction

Advancements in high-throughput sequencing technologies have played a critical role in our emerging understanding of the profound impact microbial communities have on agricultural sustainability^1,2^, diverse ecological systems^3,4^, and human health^5^. Taxonomic composition of these microbial landscapes and how they evolve within shifting ecosystems is crucial for exploring their functional roles. This characterization of microbial communities is primarily achieved through select marker gene sequencing, such as 16S ribosomal RNA (rRNA) amplicon sequencing for bacteria^6^ or Internal Transcribed Spacer (ITS) sequencing for fungi^7^. Subsequent evaluation and analysis of these sequences traditionally employs a sequence clustering approach, namely either via an Operational taxonomic unit (OTU) based approach or an Amplicon Sequence Variant (ASV) based approach. OTU-based clustering methodologies, such as Mothur^8^ and USEARCH^9^, define a taxonomic unit as a cluster of reads with a sequencing identity above some threshold, usually 97% or 99%, which reduces the overall computational complexity and mitigates the impact of sequencing errors on downstream analysis. ASV-based clustering approaches, such as DADA2^10^ and Deblur^11^, provide a similar analysis, utilizing instead 100% sequencing identity threshold followed by an error model estimation. Following clustering, taxonomic assignments are generated from classifiers such as the RDP Naïve-Bayes Classifier^12^ and MetaSquare^13^.

However, one of the challenges with these currently established methods for marker gene analysis is that they lack the sensitivity to accurately characterize strain, species, genus and even family level taxonomy. Many previous studies have shown that well-established taxonomic classifiers such as those implemented in the popular QIIME2^14^ and DADA2 workflows, are typically unable to successfully classify more than 1/3 of the reads at the species level; even when moving up the taxonomic tree to genus and family levels, these tools are still consistently unable accurately classify more than half of the reads^15-17^. This can be due to many factors, such as ambiguous 16S rRNA genes across taxonomic classification ranks^6^, potential sequencing errors, naïve clustering approaches which assume read independence, and variable 16S rRNA gene copy number^18^. However, advancements in sequencing technologies, 16S reference database curation, sequence alignment algorithms and computing performance make sequence alignment based 16S methodologies more appealing. Metagenomics-derived alignment and classification tools such as PathoScope 2.0^19^, which utilize sequence aligners such as Bowtie2^20^ and Subread^21^ have already been shown to outperform more traditional marker gene based classification approaches^22^, achieving accurate species level assignments for approximately 2/3 of the reads.

Using this rationale, we present MetaScope, a modular end-to-end R-based software package developed as an extension of the PathoScope2.0 framework. While PathoScope2.0 was originally designed as a Python-based pipeline for strain-level metagenomic and metatranscriptomic microbial abundance estimation, MetaScope reimplements these core functionalities and incorporates optimizations for both metataxonomic 16S classification and metagenomic microbial abundance estimations within an R package. In addition to the reimplementation of the maximum-likelihood read reassignment algorithm central to PathoScope, MetaScope introduces two new key features: (1) the incorporation of user-defined or empirical prior weights on candidate taxa to refine abundance estimates tailored for 16S rRNA amplicon sequences, and (2) the integration of MetaBlast, a secondary BLAST-based sequence alignment for taxonomic assignment module.

## Methods

### Implementation of Prior Weights within the PathoScope Mixture Model

To improve taxonomic resolution and reduce false-positive assignments, we implemented the PathoScopeID module in MetaScope to incorporate prior weights during the read reassignment process. The original PathoScope algorithm employs a Bayesian framework, where reads initially mapped to multiple reference genomes are reassigned probabilistically to the most likely source organism based on alignment scores and abundance estimates^23^. Specifically, we’ve implemented a user-defined or data-informed prior weights to guide the reassignment process toward biologically plausible or expected taxa. These priors can be uniform or noninformative as in the original PathoScope implementation, derived from a known sample composition such as through metagenomic sequencing, or inferred based on previously established data. Once selected, this prior is integrated into the Bayesian maximum a posteriori (MAP) estimates during the Maximization-step of the Expectation-Maximization algorithm. This is equivalent to adding a predefined number of unique reads (aj) or non-unique reads (bj) to each genome before the actual sample data is processed which biases the initial parameter estimates towards the expected composition and serves to stabilize the algorithm. Overall, this prevents estimates from converging to extreme boundary values (0 or 1), especially in scenarios characterized by sparse data.

### MetaBlast Infers Species-Level Abundance of Metataxonomic Samples by Validation of Taxonomic Assignments

Many sequence alignment-based tools in other bioinformatics fields implement multi-step or multi-alignment techniques to improve the performance and accuracy of their pipeline tools. For example, MUSCLE^24^, a tool for multiple sequencing alignment of proteins, uses a rapid alignment, followed by an improved progressive alignment, and a final refinement alignment to generate accurate multiple alignment profiles. Similarly, genome variant discovery and characterization tools such as the GATK^25^ and ABRA2^26^ use an indel realignment algorithm to increase the sensitivity in detection of certain single nucleotide polymorphisms (SNPs).

Applying the same reasoning, we’ve also developed MetaBlast, a two-step module in MetaScope for post-processing alignment and classification tool for validation of species-level taxonomic classification of amplicon-based sequencing. The first step, MetaValidate, requires an updated, standard-format BAM alignment file and a feature classification file list (containing and TaxonomyID identifier matching the reference genomes of the BAM alignment file, and read count), both of which are supplied from the MetaID function, although the module is compatible with any formal or informal alignment and classification pipelines. Reads are sampled for secondary re-alignment based on feature table prevalence and a BLAST validation matrix which consist of a probabilistic matrix for congruently aligned reads is generated to cross reference against the original read assignment. User defined validation metric cutoffs for validation metric scores such as species and genus level accuracy (i.e. the percentage of BLAST alignments that contained the original taxonomic classification), contaminant presence (i.e., the percentage of reads that either aligned or failed to align with the original MetaID assignment), and overall uniqueness (the proportion of reads whose best alignment matched the original assignment) can be applied.

Non-validated features often reflect ambiguous read alignments that may have originated from other microbes. To resolve this, a reassignment algorithm is applied to all non-validated taxa, where their BLAST alignments are cross-referenced with the MetaValidated feature table. If these reads have significant alignment scores (by default e-value < 1e-5) to other already validated features, the reads are automatically reassigned and redistributed proportionally to those validated taxa. We demonstrate that use of these proposed post-alignment steps, particularly secondary re-alignment and filtering also improves the accuracy of overall taxonomic assignment of 16S sequencing data.

### Performance Evaluation

To evaluate MetaScope as a full 16S sequence alignment and taxonomic classifier, we benchmarked its taxonomic assignment accuracy on both mock 16S microbial communities and real-world 16S rRNA datasets. We evaluated the impact of prior weights and sequence realignment on abundance estimation accuracy and assessed MetaScope’s performance relative to traditional 16S alignment tools, QIIME2 and DADA2.

### 16S amplicon sequencing analysis of mock and real-world microbial datasets

To test the validity and accuracy of our analysis pipeline, we re-analyzed publicly available mock and real-world microbial datasets. The mock data were generated by pooling equimolar concentrations genomic DNA extracted from 21 bacterial isolates, and 3 PCR primer pairs were used to amplify the V3, V4, and V5 regions^27^. 16S rRNA amplicon gene libraries were constructed and sequenced via Illlumina’s MiSeq platform. The real-world microbial data accessed from the NCBI Sequence Read Archive under BioProject accession number PRJNA242354. These data were by Botero et al. generated from respiratory tract clinical samples collected from six patients with pulmonary tuberculosis and six healthy control individuals which included nasal swabs, oropharynx swabs, and sputum^28^.

We evaluated these data using the DADA2 pipeline, QIIME2, and MetaScope. For the DADA2 analysis, all samples were filtered and trimmed using the following parameters: truncLen=c(150,150), maxN=0, maxEE=c(2,2), truncQ=2. Learned error rates were used to predict errors and perform denoising using the learnErrors() function. Paired reads were merged and chimeras were removed using removeBimeraDenova(). The silva_nr99_v138.2 RDP SILVA classifiers for species level assignment was used to classify the reads.

For the QIIME2 analysis, we used the built-in DADA2 implementation for read trimming, filtering and denoising using the default parameters. The QIIME2 silva-138-99-nb-classifier^29^ was used in feature classify and the classification output was exported.

For the MetaScope analysis, raw sequencing reads were first trimmed using Trimmomatic. The curated NCBI 16S database was used as the reference database and built into bowtie index libraries using MetaScope’s built-in mk_bowtie_index function. Reads were aligned using the built-in align_target_bowtie function with the following bowtie2 parameters: “--local -R 2 -N 0 -L 25 -i S,1,0.75 -k 25 --score-min L,0,1.88”. Priors were derived from the ground truth microbes with equal weights and varying total weights ranging from 0.5% to 200% of the total reads within each sample. Feature tables were generated with metascope_id with the convergence EM parameter set to 0.001 and the maximum iterations parameter set at 25. For the MetaBlast analysis, feature tables and updated bam files from MetaID were used for MetaValidate with the following parameters: num_results = 100, num_reads = 100, hit_list = 5. The generated validated feature table was used for MetaClean with a species_threshold = 0.5. Ground truth and accuracy metrics were calculated proportional to 16S gene copy number from rrnDB^18^.

### Software availability

MetaScope is implemented in R (≥4.5.0). The MetaBlast module interfaces with system BLASTN+ tools (≥2.12) and assumes access to appropriate reference databases. The MetaScope pipeline is available on Bioconductor (https://www.bioconductor.org/packages/release/data/experiment/html/MetaScope.html) and the development branch is available on github at: https://github.com/wejlab/MetaScope All analysis were performed in R and source code for these analyses are available in: https://github.com/seanlu96/MetaScope_16S_Benchmarking

## Results

To evaluate these 16S analysis pipelines and taxonomic classification methods, we calculated percent species-level accuracy, precision and recall metrics across a range of abundance thresholds, and microbial community distance metrics from the ground truth. Pipeline outputs were reformatted to match species taxonomy where applicable and to allow for direct comparison between classifiers. Accuracy metrics were calculated using species-level taxonomy matching from the ground truth. Reads were assigned as no species calls if they were classified at the genus level or higher within the ground truth and with no species level classification. Reads were assigned to incorrect species calls if they were classified at any taxonomic level that was not present in the corresponding taxonomic level of the ground truth.

QIIME2 and DADA2-NB on average achieved a species level with a species level accuracy of 21.1% and 34.7% respectively across all samples (Fig. 1). Additionally, QIIME2 and DADA2’s default classifiers made 5.1% and 9.3% erroneous read assignments within the mock dataset where species level or higher taxonomic level calls were not within the mock microbiome. MetaScope’s alignment and classification of these mock 16S reads improved the overall species level classification accuracy to 82.1%, and applying the optional priors and MetaBlast modules improved species accuracy to 92.9% and 93.6% respectively. Of note, MetaScope does not ambiguously assign any reads to no call as the algorithm estimates genomes of origins for these reads, and MetaScope, MetaScope with priors, and MetaBlast have an incorrect species call rate of 17.9%, 7.1% and 6.4% respectively (Fig. 1B). At the genus level, all pipelines performed well, with genus level accuracies of greather than 96% and all MetaScope pipelines achieving genus level accuracies of 99.5% (Supplemental Fig. S1B). Notably, applying priors or secondary blast reassignment did not improve genus level accuracies (Supplemental Fig. 1A).

**Fig. 1.**
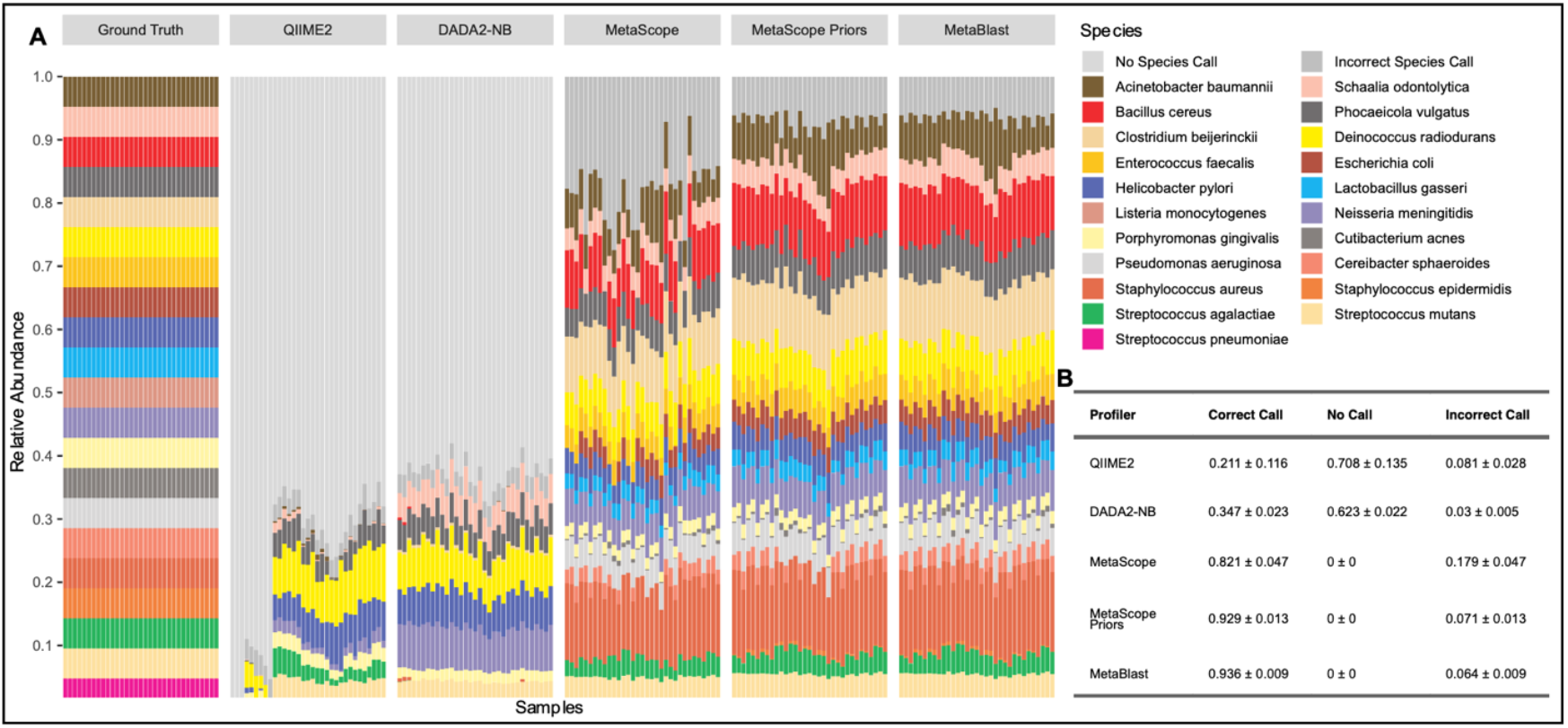
Relative abundance plots of species level accuracy for mock microbiome. **(A)** A stacked bar plot of correctly identfied microbes in the Kozich et al. mock dataset. Reads classfied in a genus or higher-level taxonomy within the ground truth are categorized as No Species Call reads and reads classfied at any taxonomic level not in the ground truth are categorized as Incorrect Species Call reads. (**B)** Summary statistics (mean and standard deviation) across 33 samples of correct call, no call and incorrect call across diHerent profilers.

To further assess the accuracy of species classification, we calculated weighted precision, weighted recall, and F1 for each pipeline. Weighted precision was determined by the proportion of correctly identified species normalized by their deviation from the true relative abundance (Fig. 2). Similarly, weighted recall was determined by the proportion of true species normalized to their deviation from the true relative abundance over the total species identified. Precision-recall (PR) curves were generated based on applying relative abundance thresholds from 0 – 0.1 at intervals of 0.0005, and F1 curves were computed as the harmonic mean between the precision and recall at each abundance threshold. Using abundance thresholds as the classification boundary, average area under the curve (AAUC) was calculated. As expected, these pipelines exhibited a tradeoff between precision and recall across the abundance thresholds. However, QIIME2 and DADA2-NB showed significantly lower AAUCs (0.089 and 0.154, respectively) compared to MetaScope and its downstream modules (MetaScope + Priors and MetaBlast), with average AAUCs ranging from 0.656 to 0.787 (Fig. 2A). In some samples, both QIIME2 and DADA2 showed a weighted precision and recall of 0: for QIIME2, this occurred for samples where no correct species were recovered at any threshold. For DADA2, these occurred at high abundance thresholds between 0.055 and 0.1 where correctly identified reads did not exceed a relative abundance. All pipelines achieved their an average maximum F1 scores across samples at low abundance thresholds, typically between 0.003 and 0.008, suggesting that many of the false positives were classified below these levels (Fig. 2B). In particular, the F1 scores without an abundance filtering (i.e. threshold of 0), were significantly lower for most samples in the QIIME2 analysis and all samples in both the DADA2 and MetaScope analyses, which highlights the prevalence of low-abundance false positives across all pipelines. As expected, F1 scores decreased when the applied abundance threshold exceeded the true relative abundances of correctly identified species (approximately 0.048) due to true positives were being excluded from the evaluation. Importantly, MetaScope with Priors and the MetaBlast modules achieved similar average maximal F1 scores of around 0.8 at abundance thresholds of 0.003.

**Fig. 2.**
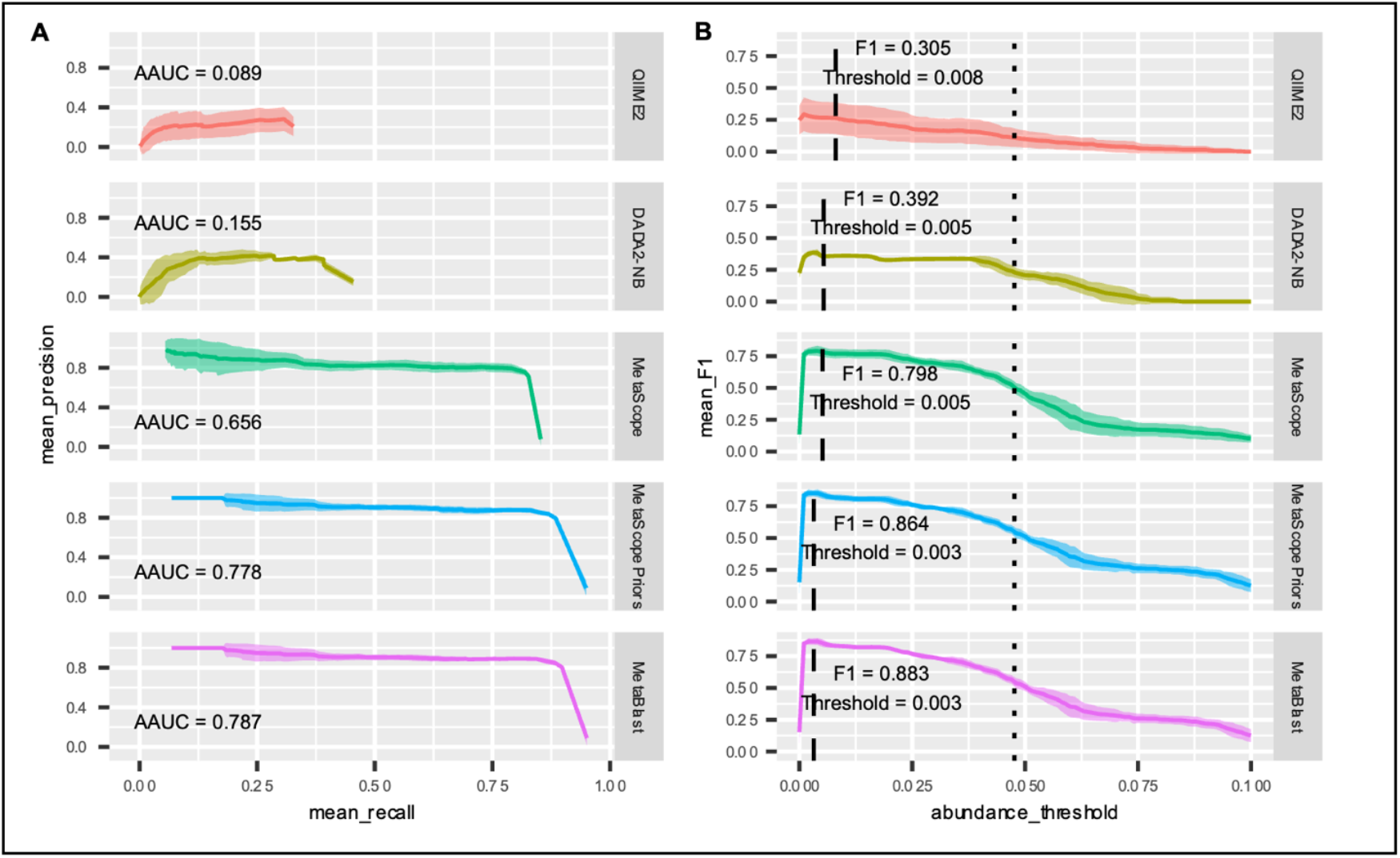
Weighted Precision-Recall Curves and F1 Curves Across Relative Abundance Thresholds. **(A)** Precision and recall were for each pipeline was calculated at relative abundance thresholds intervals of 5e-4 between 0 and 0.1 and were weighted by the deviation from the ground truth abundance. **(B)** F1 curves were calculated as the harmonic mean between weighted precision and weighted recall. Large dash lines indicate the abundance threshold where maximal F1 was achieved, and short dashed lines indicate relative abundance of all ground truth species.

**Fig. 3.**
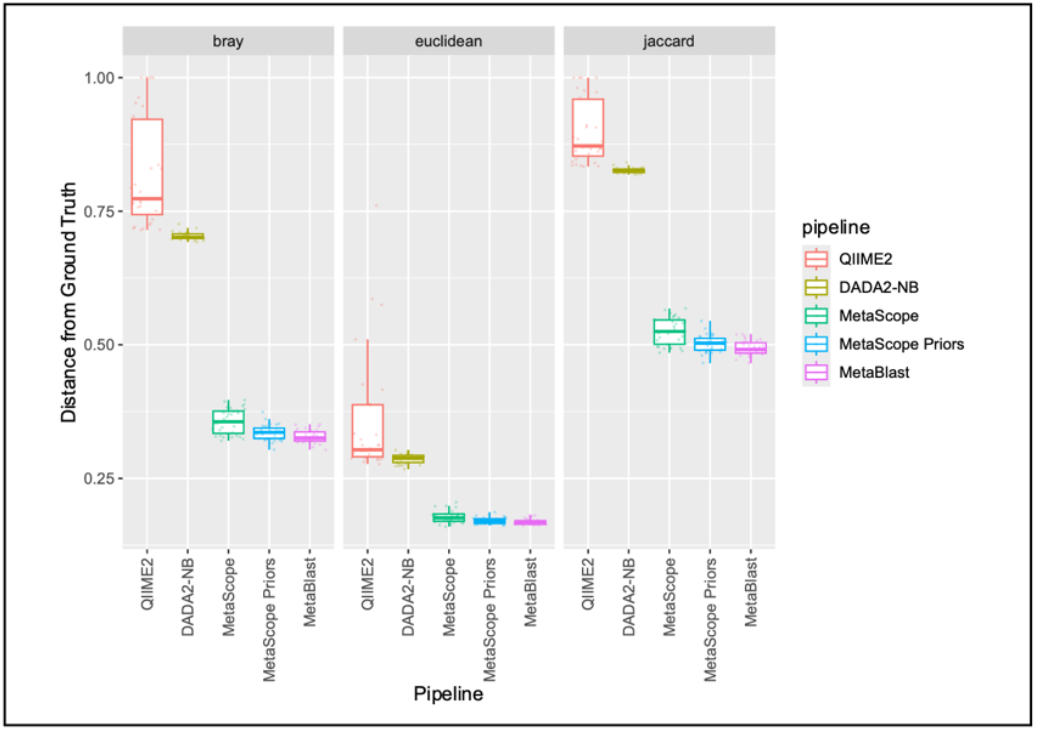
Beta Diversity Metrics from Ground Truth across pipelines. Dissimilarity indices calculated using separate three metrics (Bray-Curtis dissimilarity index, Euclidean distance, and Jaccard index) relative to the ground truth across 5 analysis pipelines.

To evaluate how species-level classification affects microbial profiling, microbial community distance metrics were calculated. Dissimilarity indices for ecological distances were calculated using Bray-Curtis dissimilarity index, Jaccard index, and Euclidean distance relative to the ground truth abundances. Similar to the relative abundance accuracy results, MetaScope and downstream modules had significantly closer Bray-Curtis, Euclidean and Jaccard distances to the ground truth species than both QIIME2 and DADA2-NB, indicating the importance of species level classification accuracy in metataxonomic analyses.

### Evaluation of MetaScope on Clinical Sequencing Samples

In addition to this mock microbiome, we evaluated MetaScope and QIIME2 and DADA2-NB on clinical samples from the Botero et. al. 16S dataset^28^. Stacked bar plots of relative species abundance of the were generated for each pipeline to compare microbial compositions across sampling sites and disease status. At the genus level both all three pipelines classify similar genera; however, QIIME2 fails to classifiy reads in the Nasal samples that both MetaScope and DADA2 classify as Staphylococcus (Supplemental Fig. 2A). At the species level, DADA2 classifies fewer reads as “uncultured bacterium” or unknown species compared to QIIME2 across all control samples and TB sputum samples. In contrast, MetaScope assigns a species-level classification to all reads, avoiding ambiguous or unclassified assignments. (Supplemental Fig. 2B). Within the nasal control samples, MetaScope identified species associated with a healthy adult nasal microbiome including *Staphylococcus aureus, Cutibacterium acnes, Corynebacterium accolens*, and *Staphylococcus epidermidis* (Fig. 4, Supplemental Fig. 2C)^30,31^. DADA2 also identified *C. accolens, C. acnes*, and *S. epidermidis*, but does not classify any of the *Streptococcus* species while QIIME2 classifies no reads at the species level with an abundance threshold of 0.05 (Supplemental Fig. 2C). In the TB nasal samples, all three tools identified *Streptococcus pneumoniae, Moraxella nonliquefaciens, and Moraxella lincolni*. MetaScope also identified *Staphylococcus Aureus, Staphylococcus caprae and Streptococcus* mitis, that both DADA2 and QIIME2 do not classify. Instead, QIIME2 and DADA2 classify *Corynebacterium pseudodiptheriticum* and *Moraxella Cattarhalis* that MetaScope does not identify (Supplemental Fig. 2C).

**Fig. 4.**
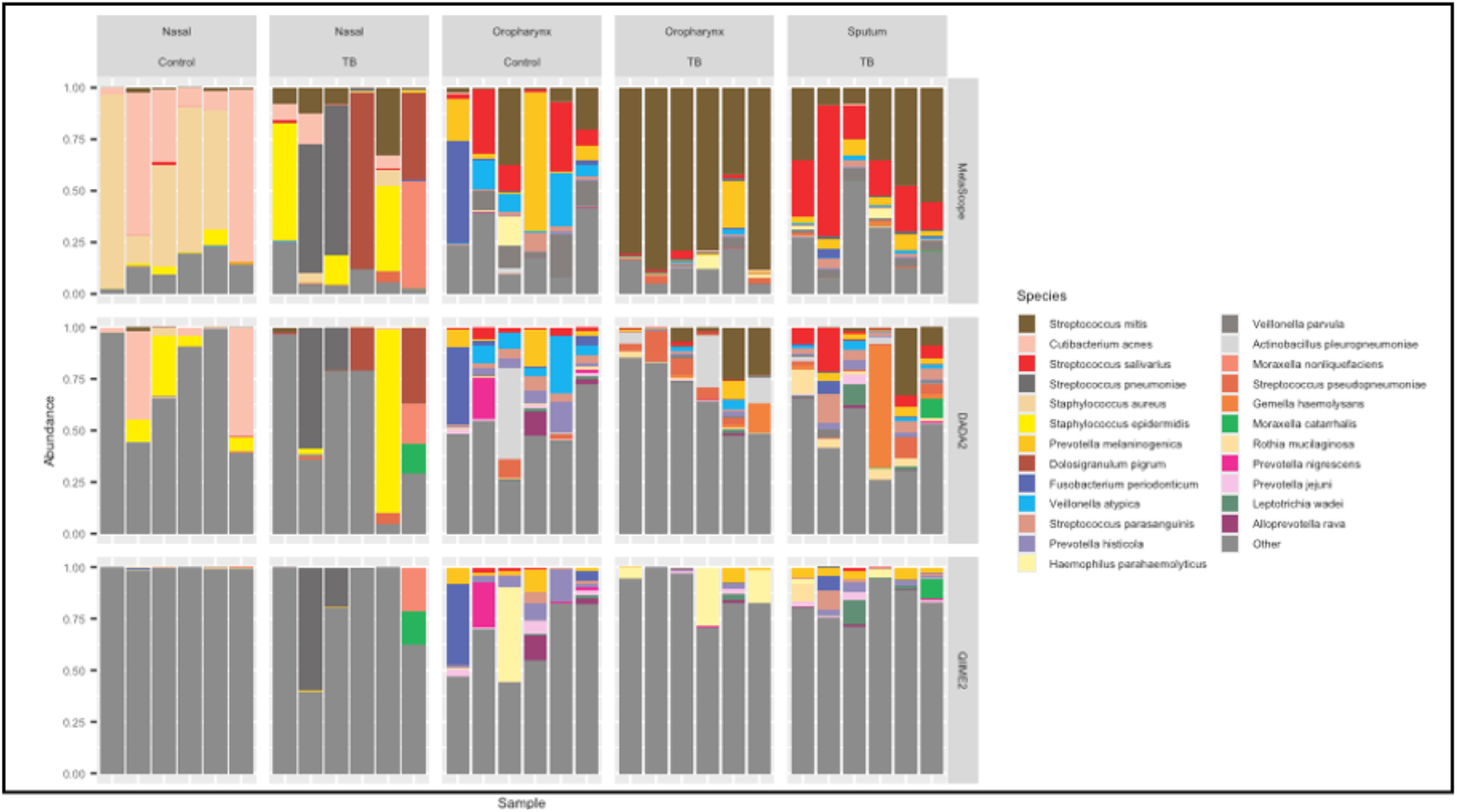
Microbial Composition of Clinical Respiratory Samples from *Tuberculosis* and Healthy Patients. Stacked bar plots of microbial species communities across healthy controls and TB patients analyzed using MetaScope, DADA2 and QIIME2. Note that no species call is grouped into other along with other low abundance microbes.

In the oropharynx samples, MetaScope identified primarily *Streptococcus salivarius, Prevotella melaninogenica*, and *Veillonella atypica* in the control samples and *Streptococcus mitis, Prevotella melaninogenica* and *Streptococcus chosunense* in the TB samples. DADA2 classifies mainly *Streptococcus vestibularis* and *Prevotella nigrescens* in the control samples and *Streptococcus mitis* in the TB samples. QIIME2 classifies *Leptotrichia*-like species and *Prevotella nigrescens* in the control samples and *Haemophilus parahaemolyticus* in the TB samples (Supplementary Fig. 3). Within the sputum of TB patients MetaScope identified primarily *Streptococcus mitis, Streptococcus salivarius*. DADA2 identifies *Gemella haemolysans* and QIIME2 classifies an unknown *Gemella* species (Supplementary Fig. 1C).

Core microbiome analysis also varied based on pipeline used. DADA2 and QIIME2 were unable to identify any core microbes at the species level using a detection rate of 0.001 and a prevalence rate of 0.2 due to a low species level sensitivity whereas MetaScope identifies 4 upper respiratory core species across all groups and the TB sputum having the most core overlapping species overall (Supplemental Fig. 2A). At the genus level MetaScope and DADA2 identify 3 core species across all groups while QIIME2 identifies 5 (Supplemental Fig. 2B).

Alpha diversity metrics between control and TB samples across nasal and oral sites showed different results based on the pipelines used. MetaScope identified a decrease in alpha diversity in between nasal control and TB samples using the Shannon, Simpson, and Inverse Simpson metrics while DADA2 and QIIME2 showed no changes in alpha diversity. In contrast, both DADA2 and QIIME2 showed a decrease in Shannon metrics but no significant changes in Simpson or Inverse Simpson in the oral samples (Figs. 5A-F).

**Fig. 5.**
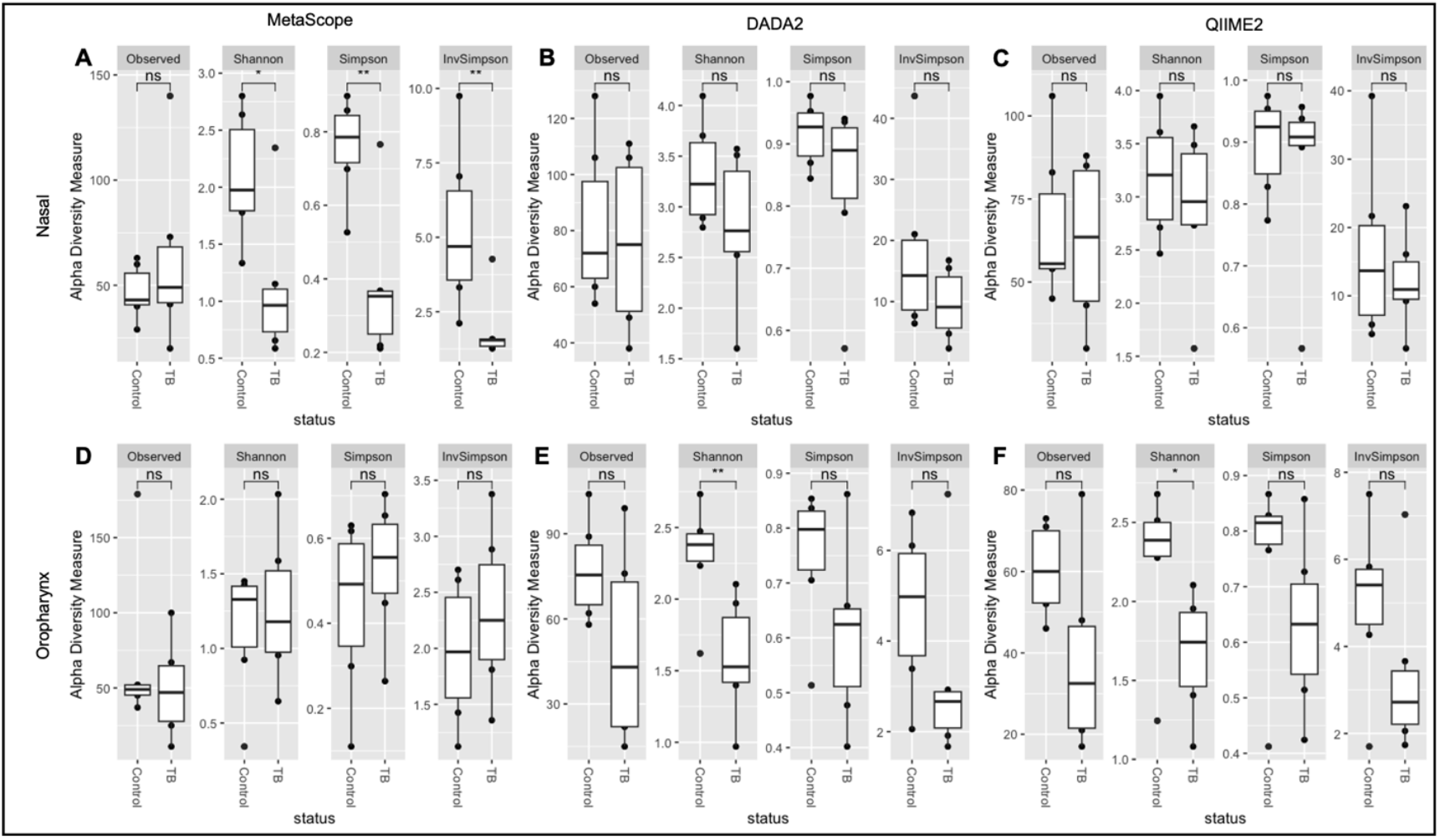
Grouped box plots of alpha diversity metrics comparing TB and control samples. (**A–C**) Nasal alpha diversity metrics; (**D–F**) Oropharyngeal alpha diversity metrics. Panels A and D: MetaScope analysis; panels B and E: DADA2 analysis; panels C and F: QIIME2 analysis.

Beta diversity analysis of the samples was performed using dimension reduction and PCoA clustering via Jensen–Shannon divergence distance metric (Fig. 6). At the genus level, all three performed similarly in clustering samples of the same site and disease status where oropharynx control and TB samples clustered closely with sputum samples and control nasal samples clustered tightly (Figs. 5D-F). At the species level, MetaScope was able to cluster the samples significantly tighter than DADA2 and QIIME2 analyses with Axis1 and Axis 2 accounting for 40.3% and 19.7% of the variance respectively (Figs. 5A-C).

**Fig. 6.**
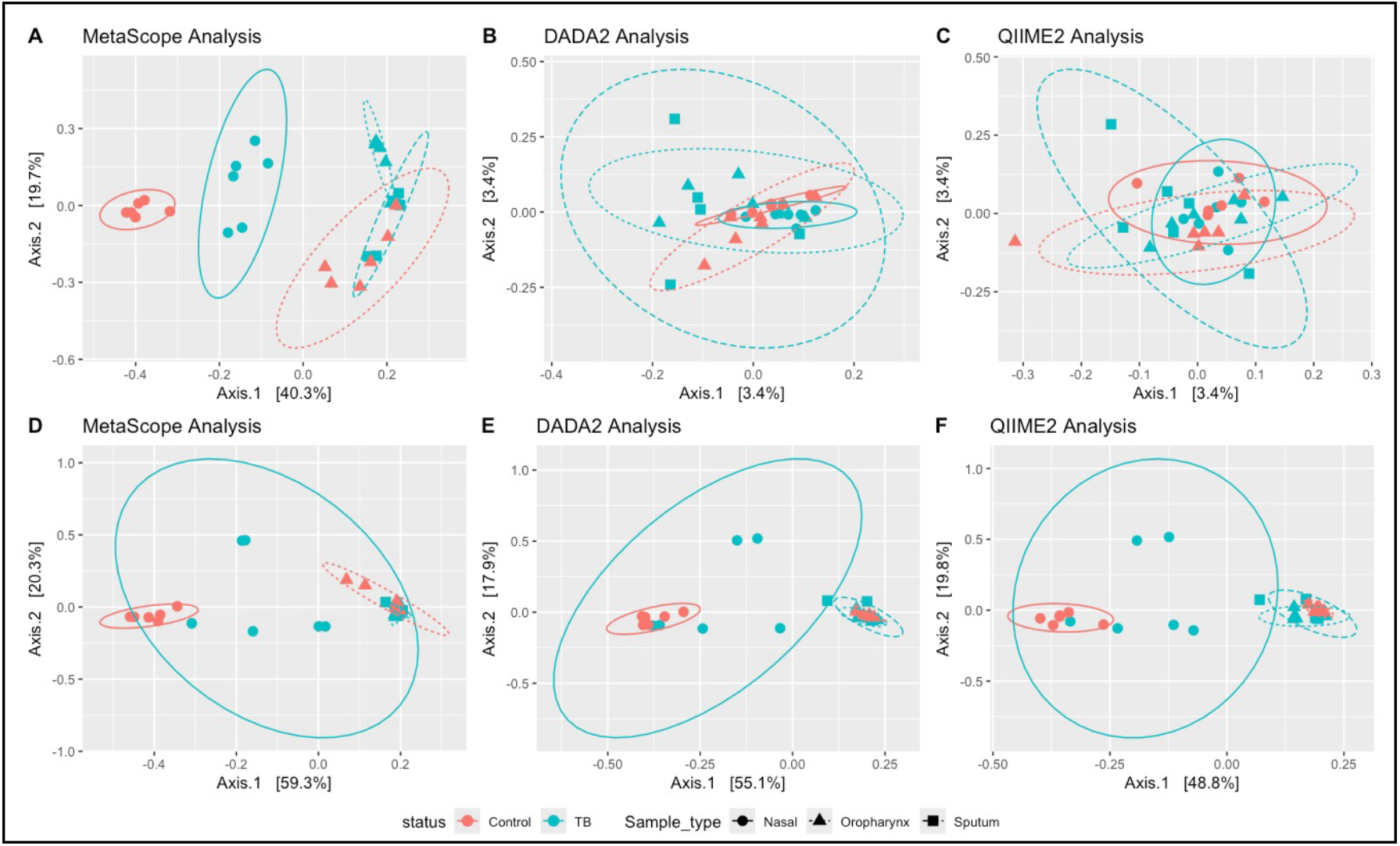
Principle Coordinate Analysis (PCoA) for TB status calculated by Jensen-Shannon Divergence Distance. (A-C) Species level Jensen Shannon Divergence Distances were used for dimension reduction. (D-E) Genus Level. Samples are clustered based on TB status (Control and TB) and sampling site (Nasal, Oropharynx and Sputum).

## Discussion

Our work underscores a paradigm shift in the classification of metataxonomic samples. Traditionally, microbial reads are clustered into Operational Taxonomic Units (OTUs) or Amplicon Sequence Variants (ASVs; e.g., via QIIME2 or DADA2), followed by taxonomic classification of these clusters. While these approaches reduce computational burden and have become standard practice, they inevitably sacrifice low-level taxonomic sensitivity and resolution. This loss of resolution not only limits microbial ecological interpretations but can also reduce statistical power in downstream analyses. In clinical contexts, inaccurate or incomplete species classification may obscure key biomarkers or pathobionts, with direct consequences for diagnostics, prognosis, and therapeutic strategies.

Advancements in sequencing technologies, alignment algorithms, reference databases, and computational resources now enable more accurate direct taxonomic classification. In this study, we present MetaScope, a taxonomy-based alignment pipeline that leverages these developments to deliver higher resolution insights into microbial diversity and community composition. By aligning reads directly against curated reference libraries, MetaScope circumvents the clustering step inherent to OTU/ASV workflows and retains the full granularity of sequence information.

Our benchmarking demonstrates that MetaScope markedly improves species-level classification in a mock microbiome, outperforming conventional 16S pipelines in both accuracy and sensitivity. In clinical samples, MetaScope was further able to identify species that remained unclassified under existing workflows, highlighting its ability to reveal microbial diversity that would otherwise remain hidden. This enhanced resolution is particularly important for the identification of low-abundance taxa, rare pathogens, or potential microbial biomarkers that may be masked by OTU/ASV clustering. Beyond improved taxonomic resolution, MetaScope also provides a framework for more consistent and reproducible microbiome profiling. Because it does not rely on arbitrary clustering thresholds, denoising heuristics, and pretrained classifiers, the pipeline reduces methodological biases and facilitates cross-study comparisons. This is especially critical as microbiome research increasingly moves toward large-scale clinical translation, where methodological consistency is paramount.

### Limitations and Future Directions

While MetaScope offers significant advantages, some limitations warrant consideration. Firstly, the accuracy of direct taxonomic alignment is dependent on the curation of reference databases; undersampled lineages or and contaminated reference databases may lead to either underrepresentation or erroneous assignments, particularly for novel taxa. Second, while computational resources have improved, alignment-based workflows can remain more computationally demanding than OTU/ASV clustering approaches, potentially limiting scalability for extremely large datasets without high-performance infrastructure. Finally, while our results demonstrate improved resolution in both mock and clinical datasets, broader validation across diverse cohorts and sequencing platforms will be essential to establish robustness and generalizability.

Nonetheless, MetaScope represents a methodological advance in metataxonomic analysis, bridging the gap between high-throughput sequencing and clinically actionable microbial insights. By enabling more precise identification of microbial community members, MetaScope enhances our ability to study ecological dynamics, discover microbial biomarkers, and interrogate the role of pathobionts in human health and disease. As sequencing datasets continue to expand in depth and breadth, we anticipate that taxonomy-based pipelines such as MetaScope will become essential tools for advancing both microbial ecology and translational microbiome research.

## Supplemental Figures

**Supplemental Fig. 1.**
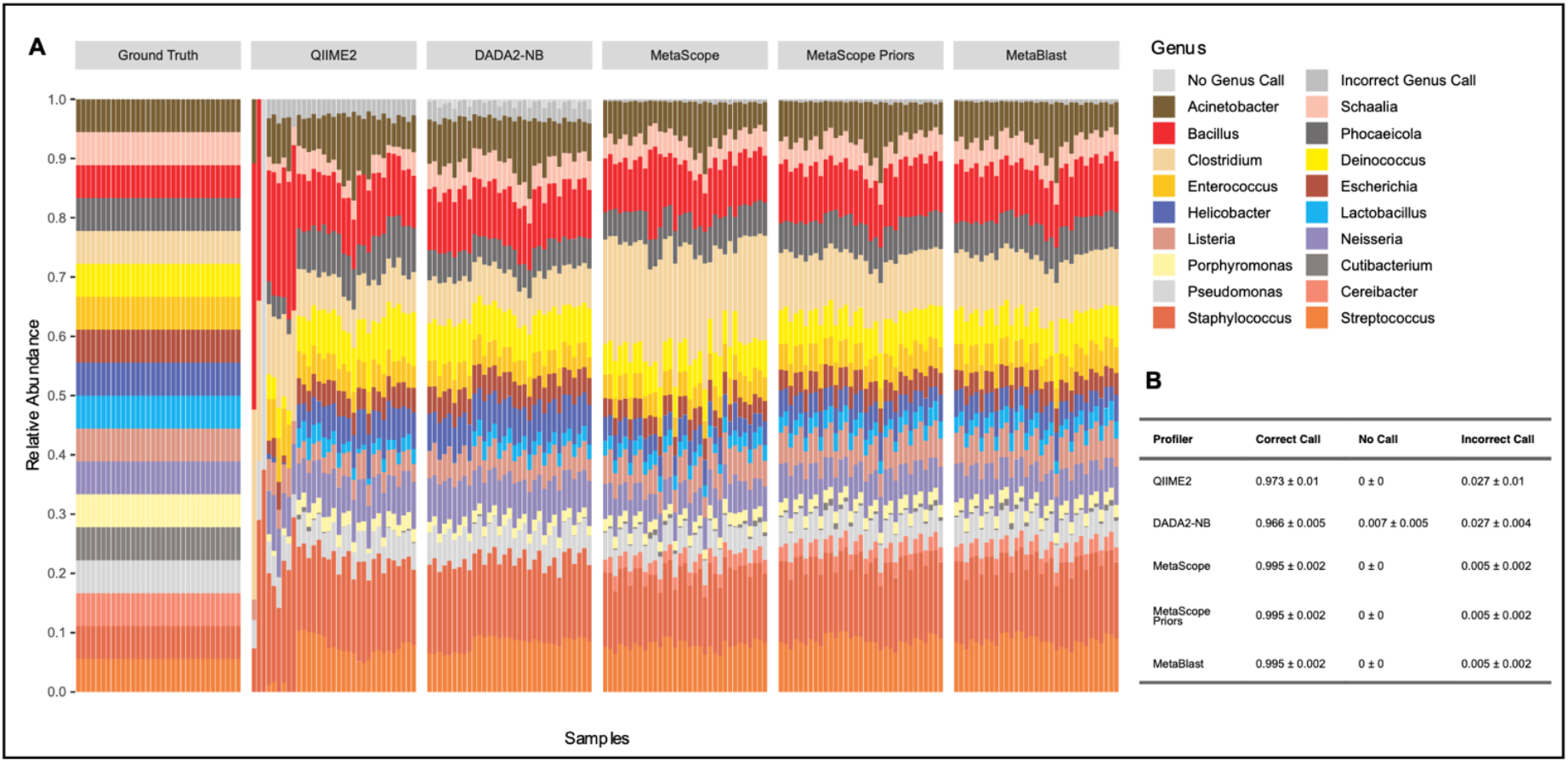
**(A)** A stacked bar plot of correctly identified microbes in the Kozich et al. mock dataset at the genus level. Reads classified in a order or higher-level taxonomy within the ground truth are categorized as No Genus Call reads and reads classified at any taxonomic level not in the ground truth are categorized as Incorrect Genus Call reads. (**B)** Summary statistics (mean and standard deviation) across 33 samples of correct call, no call and incorrect call across diHerent profilers.

**Supplemental Fig. 2.**
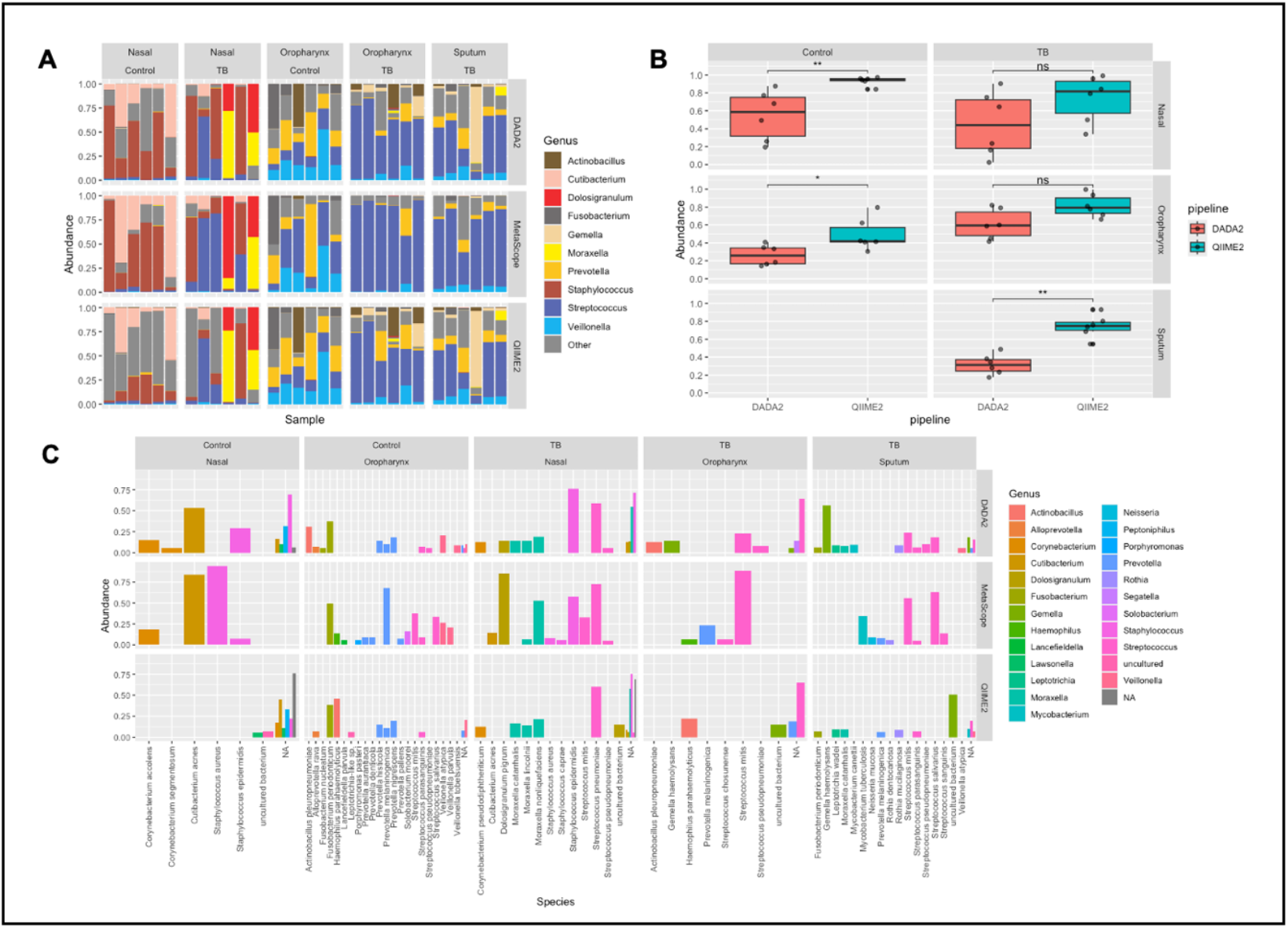
**(A)** Stacked relative abundance bar plots of microbial species communities across healthy controls and TB patients analyzed using MetaScope, DADA2 and QIIME2 grouped at genus level. **(B)** Species level abundance of “Uncultured bacterium” or “NA” no assignments for DADA2 and QIIME2. DADA2 had significantly fewer no assignments compared to QIIME2 in control samples and TB sputum samples. MetaScope had no NA or no assignment calls at the species level. **(C)** Comparison of species level assignments of all species with a relative abundance threshold greater than 0.05 in any of the three pipelines.

**Supplemental Fig. 3.**
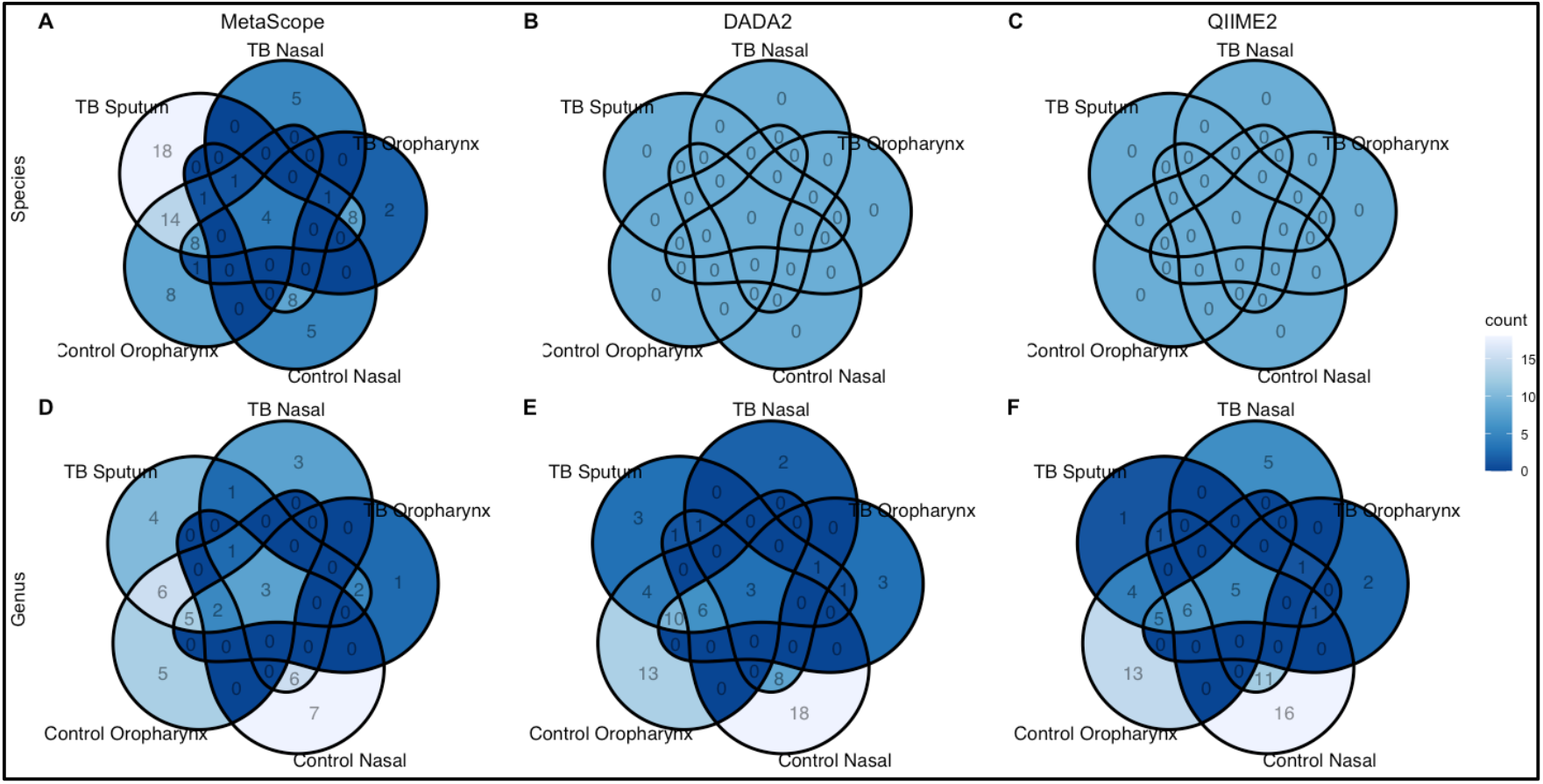
Core microbiome across all Botero et al. samples. Core microbiome analysis across all samples based on pipeline used with a detection rate of 0.001 and a prevalence of 0.2 at the species (A-C) and genus (D-F) level for MetaScope (A,D), DADA2 (B, E), and QIIME2 (C,F). Note that DADA2 and QIIME2 did not detect any core microbes at the species level due to low prevalence of species level calls.

